# Music-related Bodily Sensation Map in Individuals with Depressive Tendencies

**DOI:** 10.1101/2024.06.14.599050

**Authors:** Masaki Tanaka, Tatsuya Daikoku

## Abstract

Music has the power to influence people’s emotions. Therefore, music is also used as an intervention to reduce the stress in mental disorders such as depression and anxiety. Recent research has suggested that the body plays a key role in the connection between music and emotion with a correlation between the head sensations and negative emotions while listening to music. Additionally, strong sensations in the head have been suggested as a bodily perception associated with depression. In this study, we investigated the bodily sensations experienced by people with depressive tendencies when listening to music and their association with specific emotional states, using body mapping and musical chord progression. Our results revealed that individuals with depressive tendencies experience strong head sensations, with unpleasantness and low aesthetics, particularly for chord progression with a high level of surprise and uncertainty. This study sheds light on the intricate relationship between music, bodily sensations, and emotional states, providing valuable insights for research on the body and for developing musical therapeutic interventions targeting depression and related conditions.

## Introduction

Music is ubiquitous across human cultures^1^ and generations from infants to aged persons^2^. This is because music has the power to influence people’s emotions, for example, music can be used to calm children or to induce certain emotions in athletes before a game. Several research examined how music triggers emotional responses^3,4,5^ such as calming and stress-reducing^6^. Further, music has been used as an intervention for reducing stress in mental disorders including depression and anxiety^7,8^.

Recent studies have suggested that the body is a key to connecting music and emotion^9^. Listening to music has been shown to affect the autonomic nervous system in interaction with emotions, triggering physiological responses such as increased heart rate and breathing rate. Bodily Sensation Map (body map) has often been used as a method to visualize bodily sensations in response to music and emotions^10,11,9^. Body map is a test in which participants are asked to answer where they feel sensations in their whole body in response to stimuli, and visually demonstrates the link between emotions and the body. Specifically, it visualizes the sensations felt in specific areas of the body in response to emotions. In other words, the body map is a tool to explore where our emotions come from.

According to a previous study using body maps, there is a correlation between the head sensations and negative emotions such as anxiety and confusion, during music listening^11^. Conversely, the heart may play a strong role in the bodily perception of positive emotion (valence) associated with music^11^. Additionally, a strong sensation in the head has been suggested as a bodily perception related to depression^12^.

So, we hypothesized that when listening to music, individuals experiencing depression may have strong bodily sensations in the head, which could be associated with negative emotions. Conversely, even in cases of depression, there might be bodily sensations or musical features that evoke positive emotions. Understanding these aspects could shed light on the optimal music choices and associated bodily sensations for individuals with depression who need music intervention. However, the emotional experiences and corresponding bodily sensations during music listening in depressed individuals have not been thoroughly investigated.

Therefore, our study aims to understand how different musical structures impact individuals experiencing depression. Specifically, we examined the bodily sensations during music listening and their connection to specific emotional states. To achieve this, we conducted online experiments in body map and emotional assessments using music arranged by the eight types of four-chord progressions. These musical chord progressions consider musical uncertainty, prediction errors, and temporal dynamics, which have been shown to underlie different emotional elements in our previous studies^11^.

## Methods

### EXPERIMENTAL MODEL AND SUBJECT DETAILS

The present study consists of bodily sensation mapping tests and the following emotional judgements for every one of the eight types of 4-chord progression. We analyzed the data from our previous study^11^. The Japanese participants took part in the study (N = 527, Mean of age ± SD = ± 5.37, female = 257, specific musical training ± SD = 2.47±5.66). They had no history of neurological or audiological disorders and no absolute pitch. The experiment was conducted in accordance with the guidelines of the Declaration of Helsinki and was approved by the Ethics Committee of The University of Tokyo (Screening number: 21-335). All participants gave their informed consent and conducted the experiments by on a PC.

### METHODS DETAILS

The experimental paradigm was generated using Gorilla Experiment Builder (https://gorilla.sc), which is a cloud-based research platform that allows the deploying of behavioral experiments online. Each participant completed body-mapping tests and emotion judgements for each of the eight four-chord progressions (500ms/chord, 44.1kHz, 32bit, Electric Piano 1 based on the General MIDI, amplitude based on equal-loudness-level contours) as stimuli (Figure1.).

These stimuli were the same as those used in our previous study and were generated the 92 unique chord progressions encompassed within the eight types of chord progressions by using a statistical learning model that calculates Shannon information content and entropy based on the transition probabilities of each chord. Entropy measures the perceived uncertainty (Figure1.red line) that listeners feel in predicting the ‘next’ chord based on previous chords, while information content quantifies the surprise (Figure1.blue line) they experience when hearing the actual chord.

The four of the 8 types began with three chords, each with low uncertainty and surprise, while the other four began with three chords each displaying high uncertainty and surprise. For each set of these four progressions, the fourth chord was varying in uncertainty and surprise. Specifically, they consist of 1) sLuL-sLuL sequence (Figure 1.a) : the 1st-3rd chords have low surprise and uncertainty and the 4th chord has low surprise and uncertainty, 2) sLuL-sHuL sequence (Figure 1.b) : the 1st-3rd chords have low surprise and uncertainty and the 4th chord has high surprise and low uncertainty, 3) sLuL-sLuH sequence (Figure 1.c) : the 1st-3rd chords have low surprise and uncertainty and the 4th chord has low surprise and high uncertainty, 4) sLuL-sHuH sequence (Figure 1.d) : the 1st-3rd chords have low surprise and uncertainty and the 4th chord has high surprise and uncertainty, 5) sHuH-sLuL sequence (Figure 1.e) :the 1st-3rd chords have high surprise and uncertainty and the 4th chord has low surprise and uncertainty, 6) sHuH-sHuL sequence (Figure 1.f) : the 1st-3rd chords have high surprise and uncertainty and the 4th chord has high surprise and low uncertainty, 7) sHuH-sLuH sequence (Figure 1.g) : the 1st-3rd chords have high surprise and uncertainty and the 4th chord has low surprise and high uncertainty, and 8) sHuH-sHuH sequence (Figure 1.h) : the 1st-3rd chords have high surprise and uncertainty and the 4th chord has high surprise and uncertainty. Multiple chord progressions were generated for each of the eight types (sLuL-sLuL: 14, sLuL-sHuL: 14, sLuL-sLuH:12, sLuL-sHuH: 8, sHuH-sLuL: 18, sHuH-sHuL: 14, sHuH-sLuH: 3, sHuH-sHuH: 9), high and low thresholds were set based on the top 20% and bottom 20% of all data points for both uncertainty and surprise. The chord progression adopted was randomly selected for each participant.

**Figure 1.**
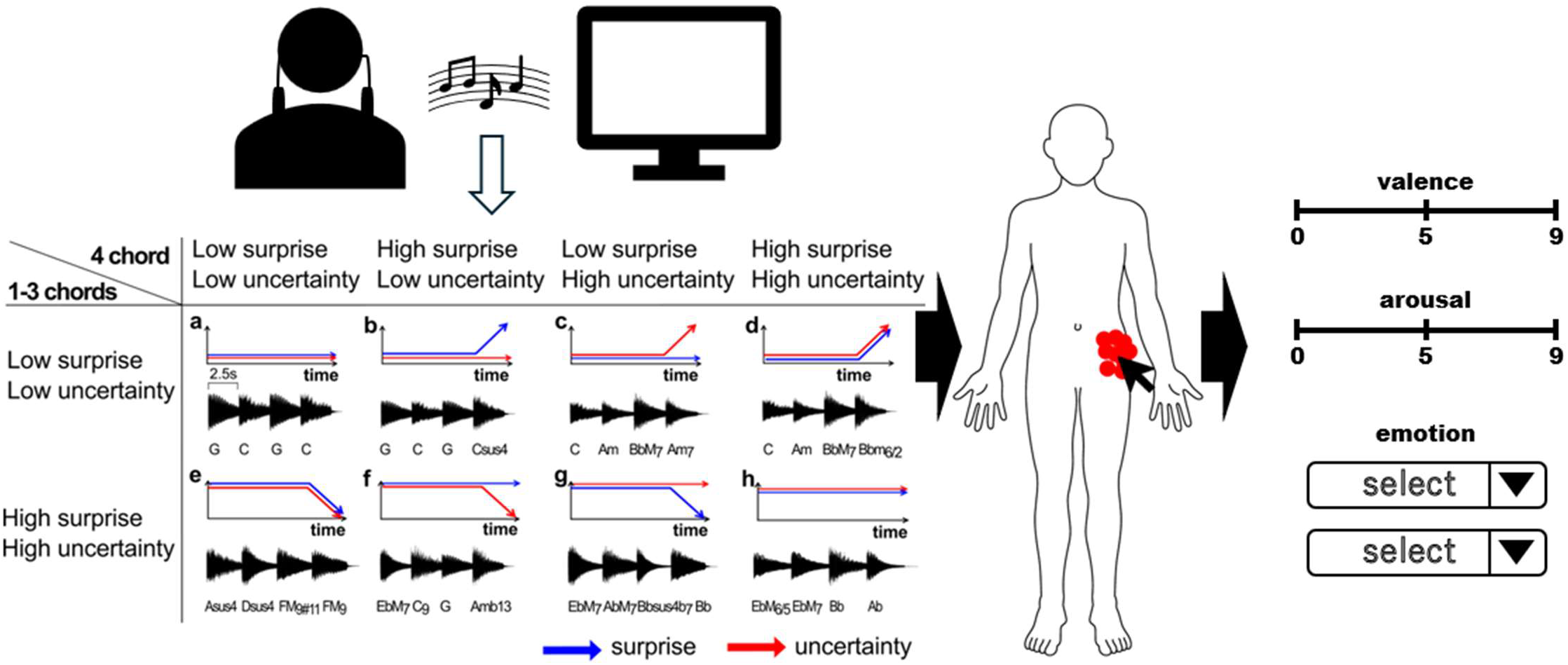
The experimental paradigm and sound stimuli. Below are 8 types of chord progressions. The blue and red arrows indicate surprise and uncertainty values, respectively.

Prior to each experiment, all participants completed the Japanese questionnaires of the Quick Inventory of Depressive Symptomatology (QIDS-J)^13,14^ (see table S1 Appendix). Using the scores from the QIDS-J, we categorized individuals into two groups according to the QIDS-J criteria. The proposed criteria are 6 to 10 for mild, 11 to 15 for moderate, 16 to 20 for severe, and 21+ for very severe depression. Thus, one group consisted of individuals with the highest scores (≧6, hereafter “depression group”), and the other group consisted of individuals with the lowest scores (≦5, hereafter “normal group”). The QIDS contains 16 items measuring the nine DSM-IV-TR criterion symptom domains, including sad mood, poor concentration, self-criticism, suicidal ideation, anhedonia, energy/fatigue, sleep disturbance, decrease/increase in appetite/weight, and psychomotor agitation/retardation, and was designed to measure the overall severity of the depressive syndrome. Each item is rated on a scale of 0-3, with higher scores indicating greater severity. The total score ranges from 0 to 27 and is calculated by summing the nine symptoms that meet the DSM-IV diagnostic criteria for major depression. The score of the items with sleep and appetite/weight is selected as the highest one among the four respectively, and psychomotor status is the same way from two. The other items are counted in their individual scores. Demographic and clinical characteristics in the Depression group are shown in Table 1.

**Table 1.**

|  | all | male | female | unknown |
| --- | --- | --- | --- | --- |
| all | 527 | 269 | 257 | 1 |
| Mean age | 32.10 | 32.60 | 31.60 | 28.0 |
| SD | 5.10 | 4.73 | 5.44 | - |
| Depression | 241 | 110 | 130 | 1 |
| Mean age | 31.73 | 32.06 | 31.48 | 28.0 |
| SD | 5.50 | 5.17 | 5.76 | - |
| Normal | 286 | 159 | 127 | - |
| Mean age | 32.42 | 32.98 | 31.72 | - |
| SD | 4.72 | 4.34 | 5.07 | - |

After each listening session in eight types of chord progressions, participants were asked to respond within 10 seconds with clicks to the position in the body where they felt sensations from the chords, using the body image presented on the screen. The clicking was allowed any number of times up to a maximum of 100 clicks, and clicking while listening to the sound was also allowed (see Figure S1 in the supplementary material for the details). In addition, each participant responded to two surveys for an emotional judgement. One was a multiple-choice categorical judgement, i.e. for each type of chord progression, participants had to select the best 5 emotion categories in the order evoked by each sound from a list of 33 categories (see Table S2 in the supplementary material). The 33 emotion categories were derived from the emotion taxonomies of prominent theorists, Keltner and Lerner^15^, and based on the previous study by Cowen et al.^16,17,18^. The other was a nine-point dimensional judgement, meaning that after listening to the chord progression, participants were asked to rate each type of chord progression along with valence and arousal. Each rating was made on a nine-point Likert scale, with the number 5 anchored in neutral.

### QUANTIFICATION AND STATISTICAL ANALYSIS

Using the coordinate data of x and y in the body mapping test, we extracted the total number of clicks at two interoceptive positions including cardiac and abdomen areas in each participant. The raw data of the x and y coordinates (see Figure S2 in the supplementary material) were down sampled by a factor of 40. The body image template used had a pixel size of 871 pixels in width and 1920 pixels in height. Specific regions were demarcated to represent distinct bodily areas: the cardiac region spanned a width from 360 to 550 pixels and a height from 390 to 620 pixels, the abdominal region was delineated between 360 to 510 pixels in width and 650 to 850 pixels in height, and the head region was specified between 240 to 601 pixels in width and 1 to 241 pixels in height. The Figures of the body topographies (Figure 2) were generated using Matlab (2022b) by interpolating the coordinates of x and y in a mesh grid format with a colour map that represented the neighboring points. The results of the best 5 emotional categories in the ranking were used to score the intensity of 33 emotions. That is, the first, second, third, fourth, and fifth categories were each scored as a 5, 4, 3, 2, and 1 point respectively. The scores of each 33 emotional categories were then averaged for all participants (see Figure S3 in the supplementary material for all results). In statistical analysis, we excluded emotions for which over 75% of responses were zero as in our previous study. As a result, 10 emotion categories including aesthetic appreciation, amusement, relief, empathy, calmness, anxiety, awkwardness, confusion, satisfaction, and nostalgia were used for further analysis.

**Figure 2.**
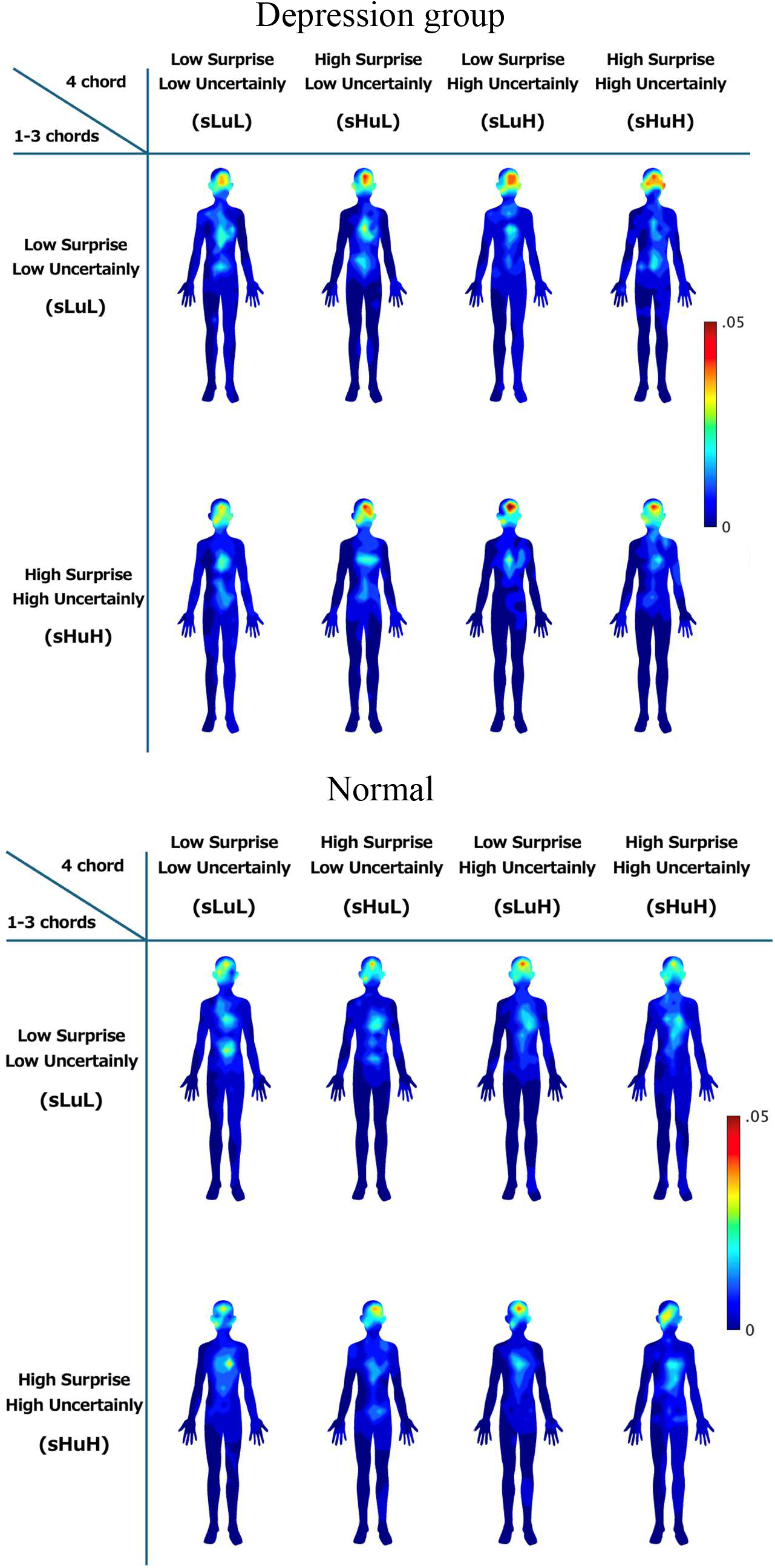
Body topography for chord progressions in the depression group. The blue-to-red gradients represent the number of clicks, ranging from few to many, respectively. In comparison to the normal group, individuals with depression show heightened sensation in the head, although the physical sensations in response to chords tended to be spread out.

We performed the Shapiro–Wilk test for normality on different click positions, the total number of clicks for each participant, the total number of clicks at cardiac, abdomen, and head positions in each participant, the 33 emotional scores of the multiple-choice categorical judgements, and the valence and arousal scores of the nine-point dimensional judgements. Depending on the result of the test for normality, the independent samples T-test and multiple comparisons were applied to compare the depression group vs. the normal group by using the data combining all chords. And then, we also performed a non-parametric (Kruskal-Wallis) One-Way analysis of variance (ANOVA) to compare the depression group to the normal group among different types of chord progressions. Further, we conducted (Spearman) correlation analysis between emotion (valence, arousal, 10 categorical emotions) and bodily sensation by using the data in the sHuH-sHuH sequence. Statistical analyses were conducted using jamovi Version 1.2 (The jamovi project, 2021). We selected p < .05 as the threshold for statistical significance and a false discovery rate (FDR) for post-hoc analysis and multiple comparisons.

## Results

We compared the topography of the codes and corresponding emotional responses in depressed and normal individuals across eight different chord progressions.

### 1. General results in listening chords in depression

Nonparametric independent sample T-tests (Mann–Whitney U test) were first conducted to ascertain differences regarding bodily sensations and feelings towards the whole chord. All the results of statistical analyses including the descriptives have been deposited to an external source (https://osf.io/m6pdt/?view_only=855bb210b9c5446592673aabd6286bf5). The results showed that individuals with depressive tendencies felt more strong physical sensations against the whole chords in the head area compared to normal individuals (*u=2.10E+06, p=.01*). In terms of emotions, the depressed individuals showed more discomfort (*u=2.10E+06, p=.03*), less aesthetics (*u=2.10E+06, p<.001*) and more nostalgia (*u=2.14E+06, p=.03*) towards the whole chords.

### 2. Specific chord to body sensation and emotion in depression

We then examined each of the individual chords mainly based on the results that were observed for the whole chords.

#### 2.1. Body map

Grand averages of the bodily map and the clicked positions in each chord progression were shown in Figures 2 and S2 in the supplemental information, respectively. All the anonymized raw data files and all the results of statistical analyses including the descriptives have been deposited to an external source (https://osf.io/m6pdt/). The individuals with depressive tendencies (Figure 2.) showed a strong sensation in the head in particular, although the physical sensations to chords tended to be diffuse. Statistical analysis indicated that the number of different click positions and the total number of clicks increased in response to the sLuL-sHuH sequence in individuals with depression group. The one-way analysis of variance (ANOVA) detected that the number of head area clicks is significantly higher in individuals with depression group compared to the normal, specifically in response to the sHuH-sHuH sequence (*χ² = 4.52, p = .03, ε² = .009*) (Figure 3.).

**Figure 3.**
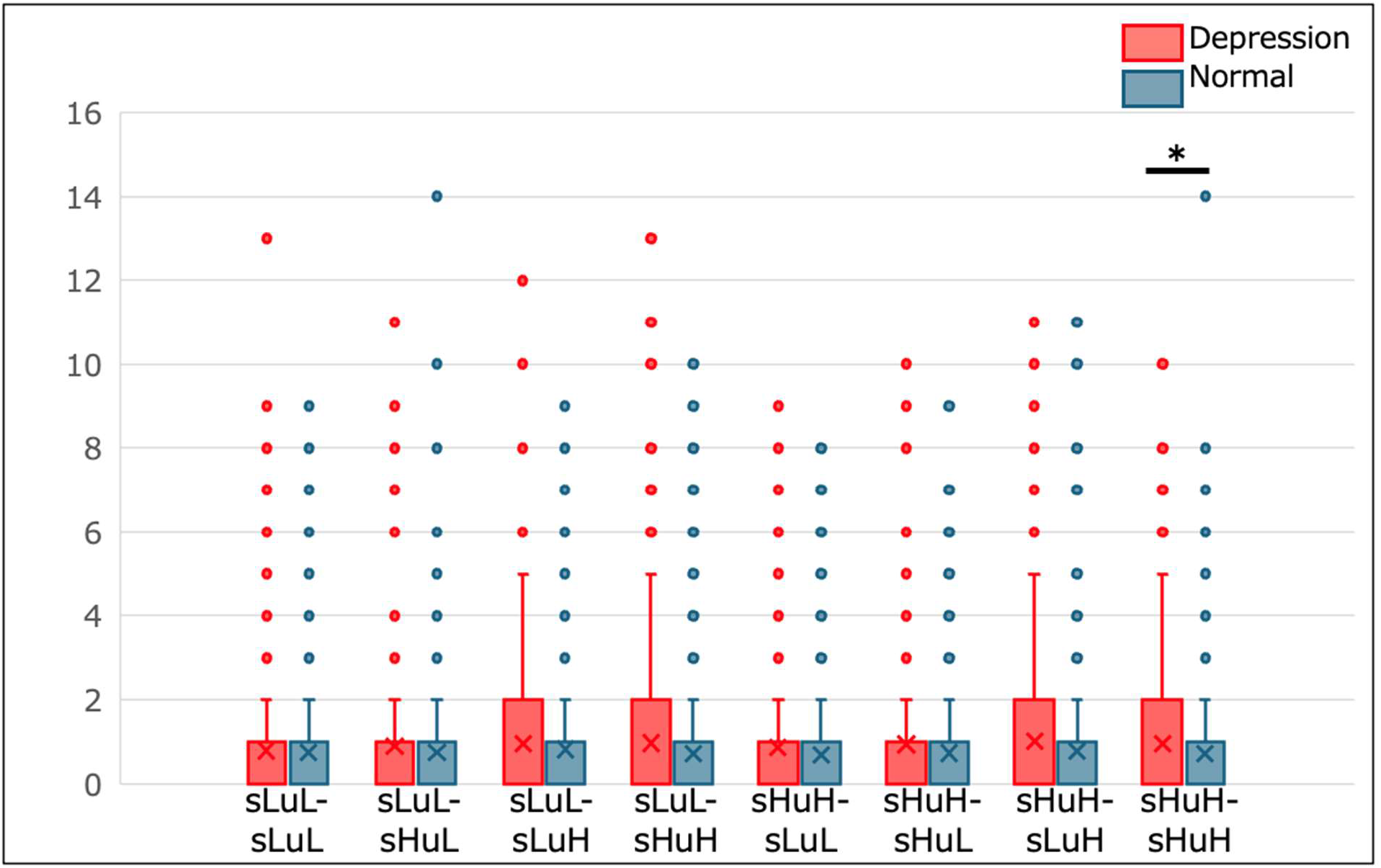
Clicks in the head area between the Depression group and the Normal. Individuals with the depression group exhibit a significantly higher number of click positions in the head compared to the normal group in response to the sHuH-sHuH sequence. \**p > .05*, One-way ANOVA.

#### 2.2. Emotion in response to chord

All the results of statistical analyses and the descriptives of the multiple-choice categorical judgements and the nine-point Likert scale of valence and arousal have been deposited to an external source. The figures of the grand average data are shown in S3 (Figure a) and S4 Appendices in supplementary materials.

The multiple-choice categorical judgements test showed that the individuals with depression felt weak aesthetics in response to several chord progressions (see Figure b, S3 Appendix). The one-way ANOVA detected that the aesthetic is significantly lower in individuals with depression group compared to the normal, specifically in response to the sLuL-sHuL sequence (*χ² = 5.08*, *p = .02*, *ε² = .01*), the sHuH-sLuL sequence (*χ² = 5.03*, *p = .03*, *ε² = .01*), and the sHuH-sHuH sequence (*χ² = 4.23*, *p < .04*, *ε² = .01*)(Figure 4.). No significant differences were found for nostalgia.

**Figure 4.**
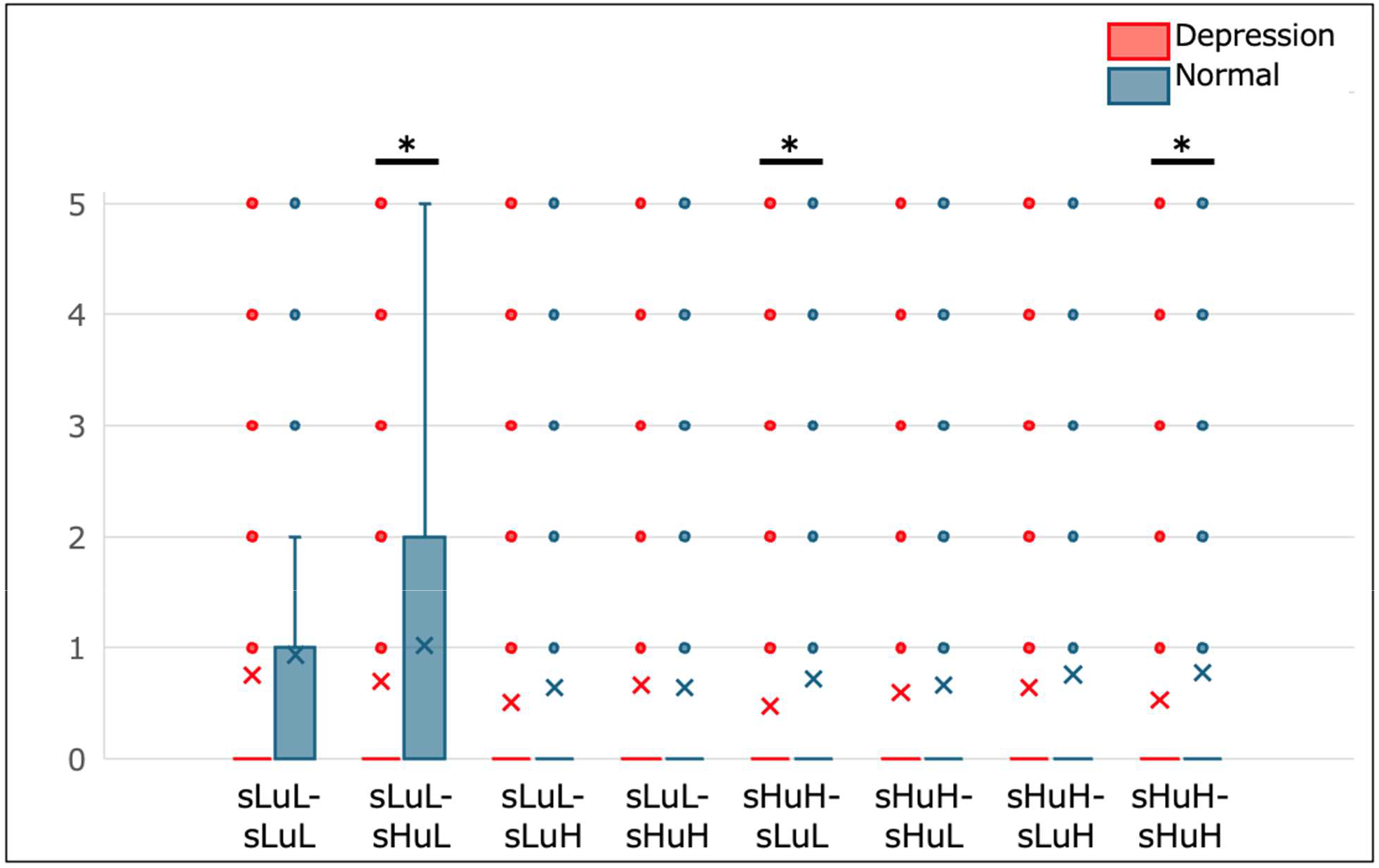
Aesthetic feeling for chord progressions in depression group. The individuals with depression felt weak aesthetic in response to several chord progressions \**p > .05*, One-way ANOVA.

The nine-point Likert scale of valence and arousal detected that the individuals in the depression group felt negative valence, while significant differences in arousal between the individuals in the depression group and the normal were not observed (Figure 5). The one-way ANOVA detected that the valence is significantly negative in individuals with depression group compared to the normal, in response to the sHuH-sHuH sequence (*χ² = 4.36*, *p = .04*, *ε² = .01*) (Figure 5.).

**Figure 5.**
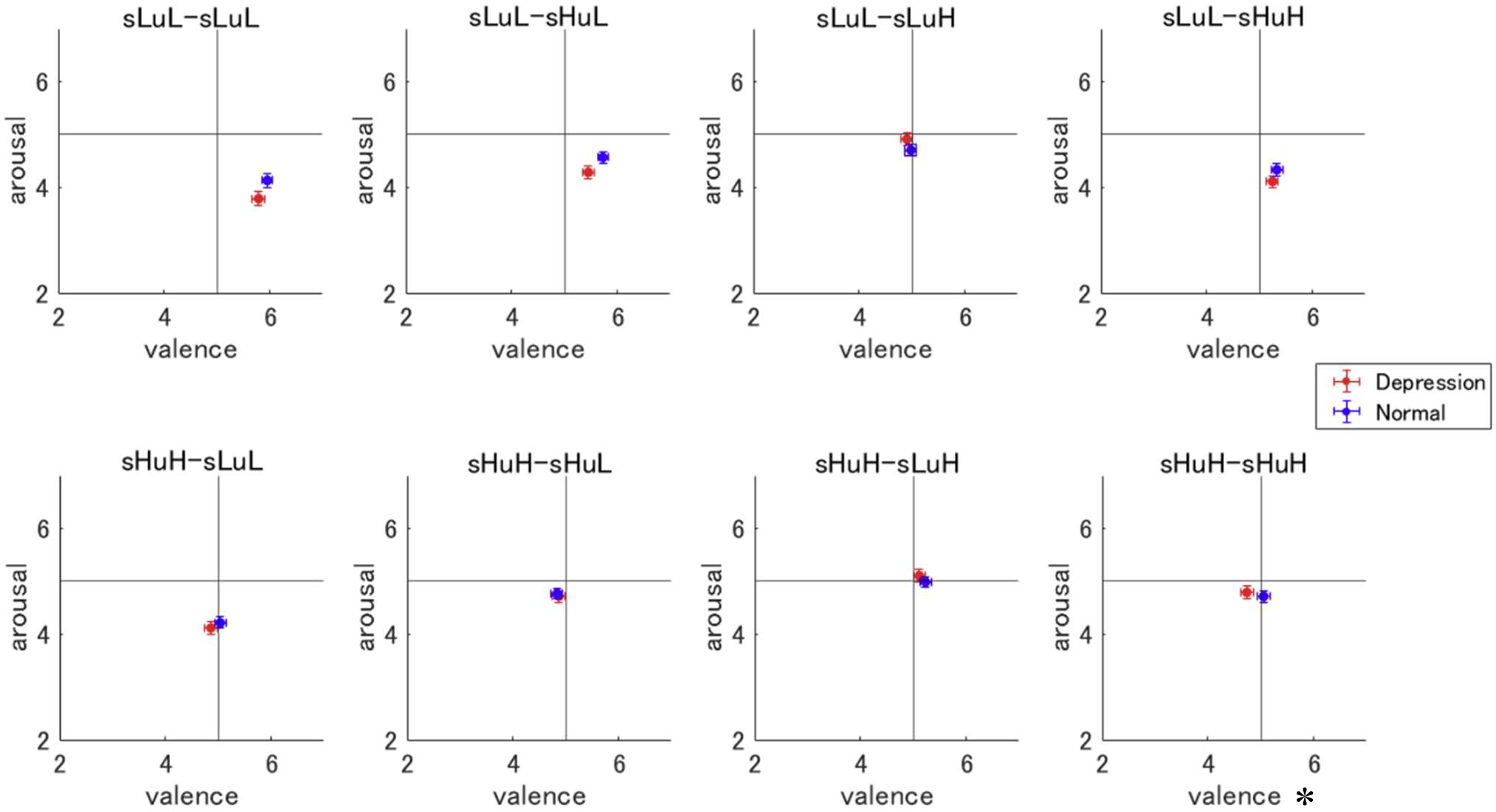
Valence and Arousal for chord progressions in depression group. The valence was significantly negative in individuals with depression group compared to the normal, in response to the sHuH-sHuH sequence. \**p > .05*, One-way ANOVA.

### 3. Emotions related to head sensations in the sHuH-sHuH sequence

Individuals with depressive tendencies showed more head or negative emotion in the sHuH-sHuH chord progression in both body map and emotional judgements compared to normal individuals. Therefore, we checked the emotions associated with the head in the sHuH-sHuH sequence in the depression group. We applied the Spearman correlation tests, as the Shapiro–Wilk test for normality showed the violations of the assumption of normality on all the data (*p < .001*). Results revealed significant positive correlations between the anxiety with the number of clicks localized to the head area (*rs = 0.178, p = .006*), although no correlation with the score of QIDS-J was found. This may suggest that head sensation is an important factor for negative emotion with depression tendencies regardless of depressive symptoms in this type of chord progression.

## Discussion

In this study, we hypothesized that individuals with depression might experience strong head sensations when listening to music, which could be associated with negative emotions. The results supported the hypothesis, revealing that those with depressive tendencies felt more intense physical sensations in the head area in response to whole chords compared to non-depressed individuals. Further, the depressed individuals exhibited greater discomfort, lower aesthetic appreciation, and increased nostalgia towards the musical chords. Specifically, in the depression group, strong head sensations, heightened discomfort, and diminished aesthetic emotions were notably associated with the sHuH-sHuH chord progression.

In our previous study^11^ we used a similar paradigm to reveal the relationship between musical perception, interoception and the resulting emotional experiences. In the current study, we provide further insight into these relationships, considering individual differences in depressive tendencies. The previous studies showed that the sLuL-sLuL and sLuL-sHuL sequences elicited predominantly abdominal and heart interoceptive sensations, and suggested a robust relationship between heart sensation and pleasant feelings. Head sensations were also significantly correlated with certain negative emotions such as anxiety and confusion. In this study, although heart and valence, relief, satisfaction, nostalgia, and calmness were positively correlated with all chords even in depressive states, individuals with depressive tendencies experienced stronger head sensations and indicated more discomfort and less aesthetic compared to control participants in the sHuH-sHuH chord progression. Head sensations were also significantly correlated with anxiety in whole chord progression like previous study. These results may indicate that head sensation is one of the keys to understanding generating negative sensations when listening to music, especially in individuals with depressive tendencies.

The past study by Sakka and Juslin^19^ indicated that depressed people are characterized by negative cognitive tendencies, called cognitive biases, which are observed in at least four areas of information processing. The four are 1)interpretation bias^20^, 2) negative attention bias^21,22,20,23^, 3) impaired in perceiving and recognizing emotions with music^24,25^, and 4) Mood-congruent memory in relation to retrieval of autobiographical (episodic) memories^26,27^ and overgeneral autobiographical memory^28^. Of note is the interpretation bias, where people with depression have been shown to have a negative bias when interpreting ambiguous events, including events that involve surprise^29^. In our results, depressed individuals showed negative affect in the sHuH-sHuH chord progression, a chord with high prediction error and uncertainty up to the third chord, which remained unstable through the fourth chord. Thus, the negative feelings towards certain chords expressed by the depressed tendencies in the present study may lead to a bias in the interpretation of certain chords when depressed people listen to music, and the fact that the results were obtained in an everyday life rather than a laboratory setting may provide new support for previous research. And the effects of music appreciation on mental health are not always positive, and in some cases, unhealthy music appreciation behaviors, such as maladaptive emotional regulation by music, have been shown to promote psychopathology^30^, so it will hope that this study will contribute to the advancement of music therapy and its research.

As for the physical sensation of emotion with music, there have been many studies on heartbeat and breathing sensations^31^, but we are not aware of any studies to understand the factors that contribute to the physical sensation in the head. According to past literature, brain waves from listening to positive and negative music show that people think more about cognitive issues when they are in a negative state compared to when they are in a positive state, leading to cortical desynchronization^32^. Although nostalgia was also reported in the current results, previous studies have suggested that although depressed individuals experience nostalgia with music, they do not necessarily benefit from nostalgic memories with music and that nostalgic memories may lead to negative emotional experiences for these individuals^33,34^. Thus, depressed people may be more prone to negative states. Furthermore, earworms are known for the relationship between the head and music, and research is being conducted to elucidate the mechanism. According to Arthur^35^, in several studies^36,37,38^, neurotic tendencies such as depression have been shown to be strongly related to ruminative tendencies and positively related to earworm experiences. Thus, Arthur insisted on the possibility that people with anxiety and nervous temperaments tend to habitually ruminate, and as a result, are more likely to develop earwigs. However, our current study utilized only the body map and emotion questionnaire and did not examine physiological approaches or earworms, so we were unable to validate these previous literatures or elucidate why individuals with depression tendencies experience strong physical sensations in their heads when they listen to music. As a next step, we will investigate multimodal methods, including physiological measurements, to elucidate the mechanism of physical sensations in the head.

In summary, this study found that depressed people experience strong head sensations and negative feelings when listening to certain musical chord sequences. Conversely, our previous research indicated that some chord progressions evoke positive interoceptive sensations in the heart and abdomen. it is crucial to understand how people emotionally experience music in their daily lives from both positive and negative perspectives, considering individual differences. This understanding could elucidate how music exerts both physical and mental influences, enhancing its therapeutic potential.

## Data Availability

All anonymized raw data files, stimuli used in this study and the results of statistical analysis have been deposited to an external source (https://osf.io/m6pdt/?view_only=855bb210b9c5446592673aabd6286bf5). The other data are shown in supplementary data.

## Acknowledgements

This research was supported by the Japan Society for the Promotion of Science (JSPS), Fostering Joint International Research (22KK0157) Japan. The funding sources had no role in the decision to publish or prepare the manuscript.

## Author Contributions Statement

M.T. and T.D. conceived the experimental paradigm and method of data analysis. M.T. collected the data, analysed the data, and wrote the draft of the manuscript and figure. M.T. and T.D. edited and finalized the manuscript.

