## Supplementary for "Music-related Bodily Sensation Map in Individuals with Depressive Tendencies"

Figure S1. Body image and example of the clicks to the position in the body.

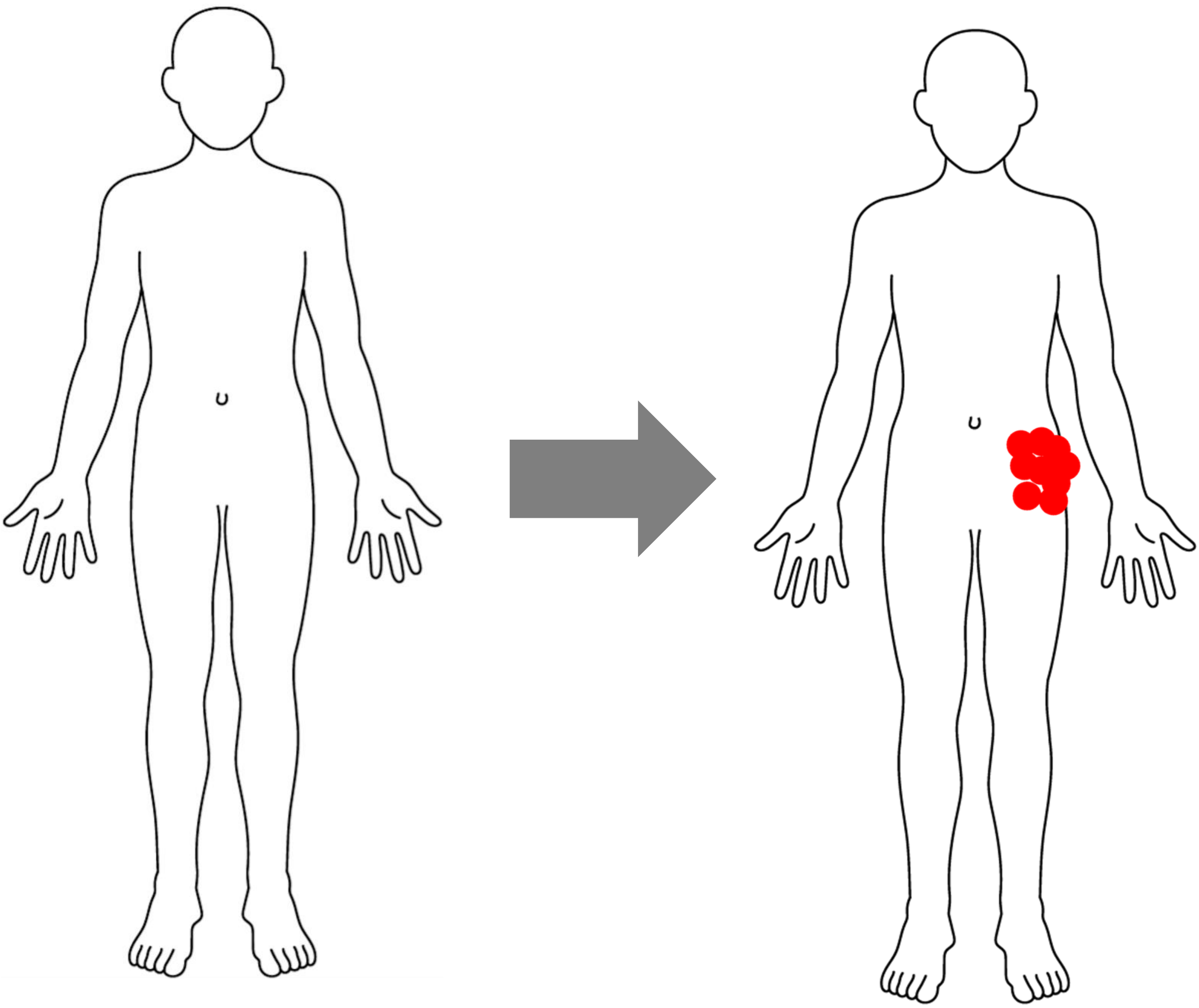

Figure S2. The row data of the click of the body.

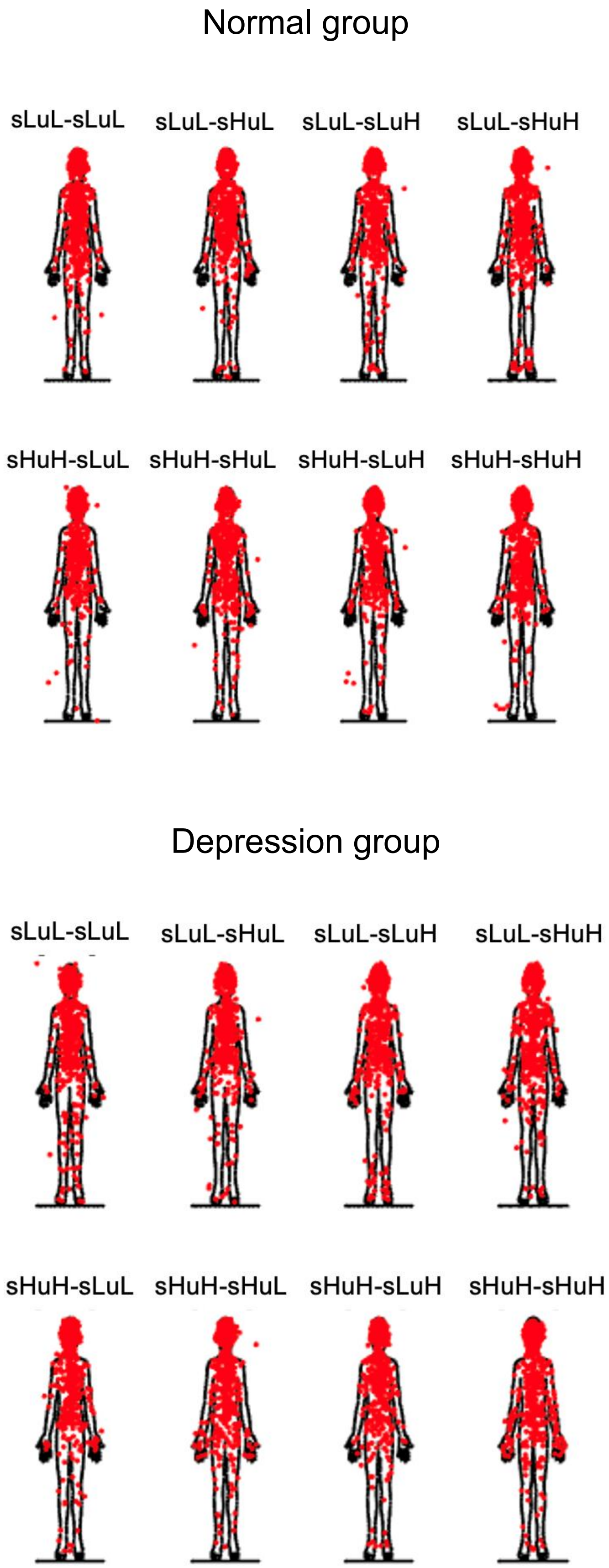

Figure S3. Categorical judgement of 33 emotions

a. all participants

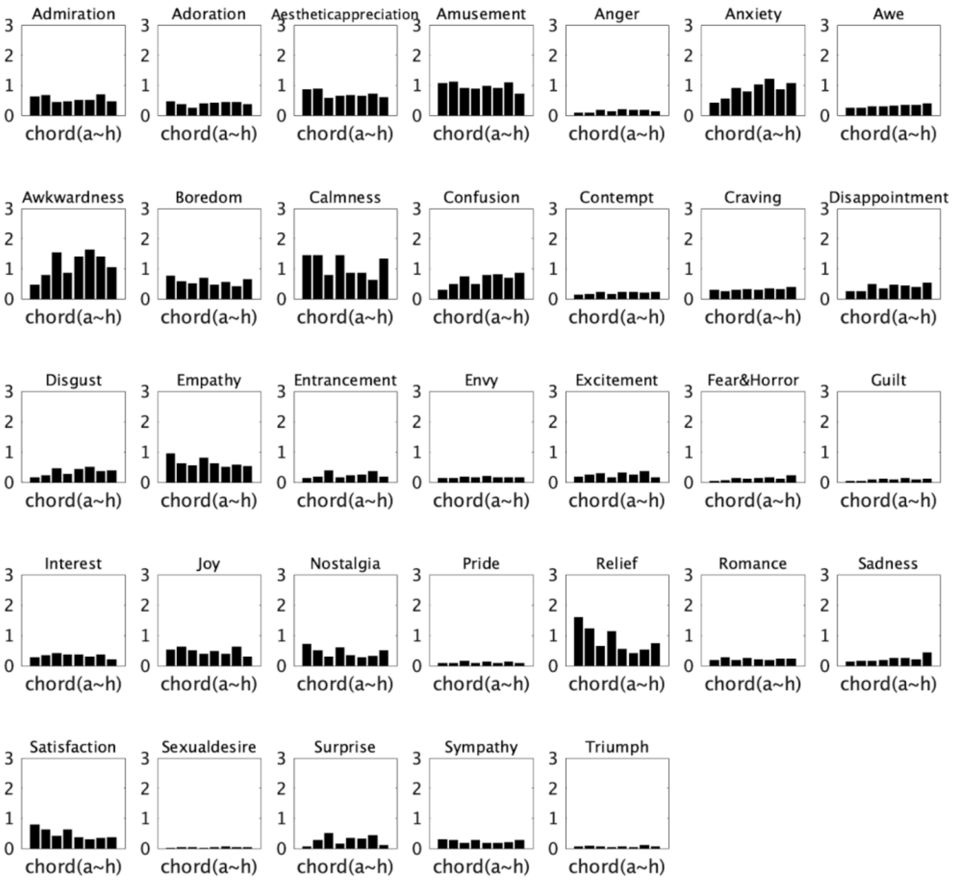


b. depression group vs normal group

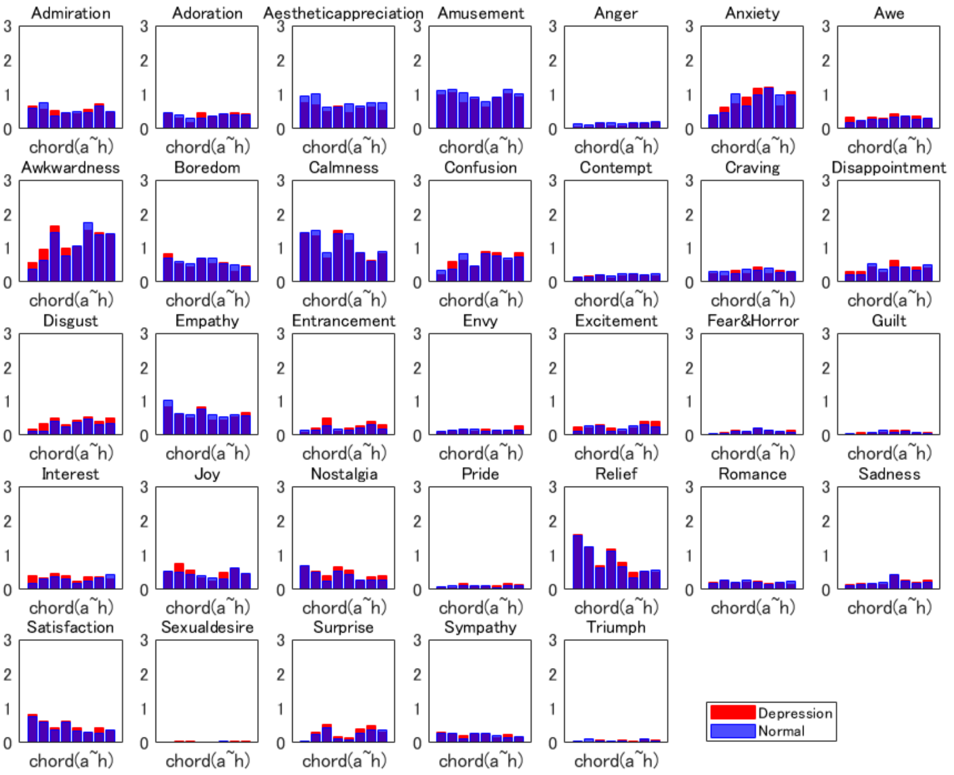

Figure S4. Valence and arousal for chord progression

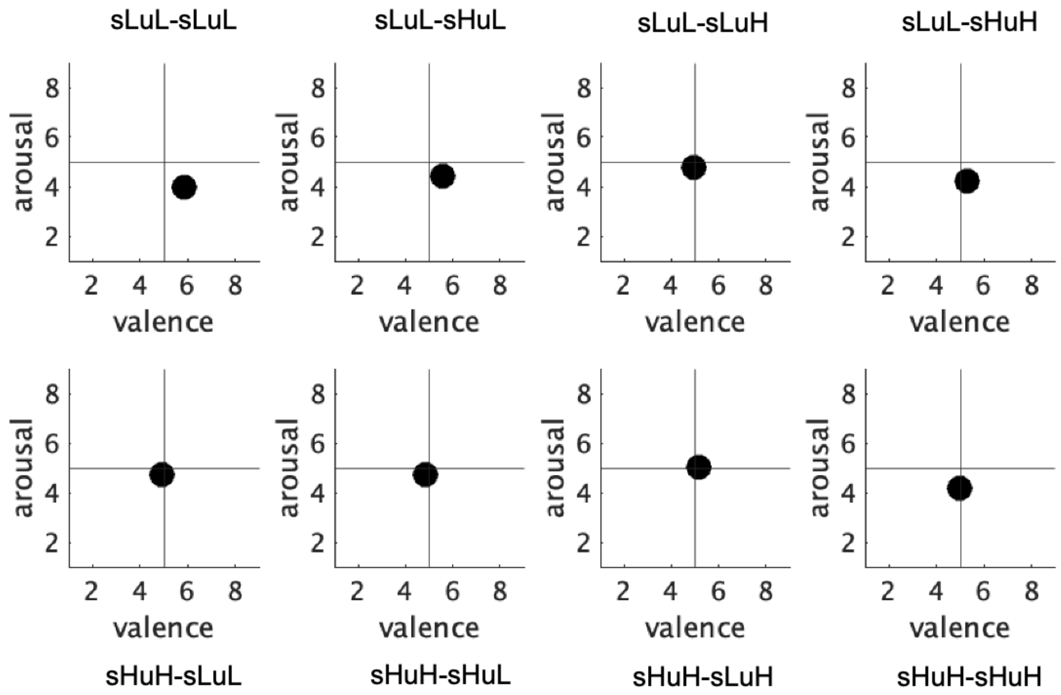

Table S1. Quick Inventory of Depressive Symptomatology

QIDS-J (Japanese version) : you can access from here.

1. Falling Asleep:

    0 I never take longer than 30 minutes to fall asleep.

    1 I take at least 30 minutes to fall asleep, less than half the time.

    2 I take at least 30 minutes to fall asleep, more than half the time.

    3 I take more than 60 minutes to fall asleep, more than half the time.

2. Sleep During the Night:

    0 I do not wake up at night.

    1 I have a restless, light sleep with a few brief awakenings each night.

    2 I wake up at least once a night, but I go back to sleep easily.

    3 I awaken more than once a night and stay awake for 20 minutes or more, more than half the time.

3. Waking Up Too Early:

    0 Most of the time, I awaken no more than 30 minutes before I need to get up.

    1 More than half the time, I awaken more than 30 minutes before I need to get up.

    2 I almost always awaken at least one hour or so before I need to, but I go back to sleep eventually.

    3 I awaken at least one hour before I need to, and can't go back to sleep.

4. Sleeping Too Much:

    0 I sleep no longer than 7-8 hours/night, without napping during the day.

    1 I sleep no longer than 10 hours in a 24- hour period including naps.

    2 I sleep no longer than 12 hours in a 24- hour period including naps.

    3 I sleep longer than 12 hours in a 24-hour period including naps.

5. Feeling Sad:

    0 I do not feel sad.

    1 I feel sad less than half the time.

    2 I feel sad more than half the time.

    3 I feel sad nearly all of the time.

6. Decreased Appetite:

    0 There is no change in my usual appetite.

1 I eat somewhat less often or lesser amounts of food than usual.

2 I eat much less than usual and only with personal effort.

3 I rarely eat within a 24-hour period, and only with extreme personal effort or when others persuade me to eat.

7. Increased Appetite:

0 There is no change from my usual appetite.

1 I feel a need to eat more frequently than usual.

2 I regularly eat more often and/or greater amounts of food than usual.

3 I feel driven to overeat both at mealtime and between meals.

8. Decreased Weight (Within the Last Two Weeks):

0 I have not had a change in my weight.

1 I feel as if I've had a slight weight loss.

2 I have lost 2 pounds or more.

3 I have lost 5 pounds or more.

9. Increased Weight (Within the Last Two Weeks):

0 I have not had a change in my weight.

1 I feel as if I've had a slight weight gain.

2 I have gained 2 pounds or more.

3 I have gained 5 pounds or more.

10. Concentration/Decision Making:

0 There is no change in my usual capacity to concentrate or make decisions.

1 I occasionally feel indecisive or find that my attention wanders.

2 Most of the time, I struggle to focus my attention or to make decisions.

3 I cannot concentrate well enough to read or cannot make even minor decisions.

11. View of Myself:

0 I see myself as equally worthwhile and deserving as other people.

1 I am more self-blaming than usual.

2 I largely believe that I cause problems for others.

3 I think almost constantly about major and minor defects in myself.

12. Thoughts of Death or Suicide:

 0 I do not think of suicide or death.

 1 I feel that life is empty or wonder if it's worth living.

 2 I think of suicide or death several times a week for several minutes.

 3 I think of suicide or death several times a day in some detail, or I have made specific plans for suicide or have actually tried to take my life.

13. General Interest:

 0 There is no change from usual in how interested I am in other people or activities.

 1 I notice that I am less interested in people or activities.

 2 I find I have interest in only one or two of my formerly pursued activities.

 3 I have virtually no interest in formerly pursued activities.

14. Energy Level:

 0 There is no change in my usual level of energy.

 1 I get tired more easily than usual.

 2 I have to make a big effort to start or finish my usual daily activities (for example, shopping, homework, cooking or going to work).

 3 I really cannot carry out most of my usual daily activities because I just don't have the energy.

15. Feeling slowed down:

 0 I think, speak, and move at my usual rate of speed.

 1 I find that my thinking is slowed down or my voice sounds dull or flat.

 2 It takes me several seconds to respond to most questions and I'm sure my thinking is slowed.

 3 I am often unable to respond to questions without extreme effort.

16. Feeling restless:

 0 I do not feel restless.

 1 I'm often fidgety, wringing my hands, or need to shift how I am sitting.

 2 I have impulses to move about and am quite restless.

 3 At times, I am unable to stay seated and need to pace around.

Table S2. 33 categories of emotion

| | |
|---|---|
| 1 | Awkwardness |
| 2 | Envy |
| 3 | Pride |
| 4 | Romance |
| 5 | Anxiety |
| 6 | Empathy |
| 7 | Craving |
| 8 | Triumph |
| 9 | Sympathy |
| 10 | Joy |
| 11 | Confusion |
| 12 | Disappointment |
| 13 | Amusement |
| 14 | Disgust |
| 15 | Relief |
| 16 | Adoration |
| 17 | Anger |
| 18 | Sexual desire |
| 19 | Fear/Horror |
| 20 | Sadness |
| 21 | Nostalgia |
| 22 | Satisfaction |
| 23 | Entrancement |
| 24 | Awe |
| 25 | Admiration |
| 26 | Guilt |
| 27 | Aesthetic appreciation |
| 28 | Interest |
| 29 | Excitement |
| 30 | Contempt |
| 31 | Boredom |
| 32 | Calmness |
| 33 | Surprise |
